# A genetically-encoded nanobody sensor reveals conformational diversity in β-arrestins orchestrated by distinct seven transmembrane receptors

**DOI:** 10.1101/2024.02.21.581355

**Authors:** Parishmita Sarma, Vendula Nagy Marková, Nashrah Zaidi, Annu Dalal, Sudha Mishra, Manish K. Yadav, Gargi Mahajan, Nabarun Roy, Paul Miclea, Josef Lazar, Arun K. Shukla

**Affiliations:** Department of Biological Sciences and Bioengineering, Indian Institute of Technology, Kanpur 208016, India; First Faculty of Medicine, Charles University in Prague, Kateřinská 32, 121 08 Prague; Institute of Organic Chemistry and Biochemistry CAS, Flemingovo nám. 2, 160 00 Prague; Faculty of Science, Charles University in Prague, Albertov 6, 128 00 Prague, Czech Republic

**Keywords:** GPCRs, β-arrestins, cellular signaling, biosensors, nanobodies, intrabodies, polarization microscopy, linear dichroism

## Abstract

Agonist-induced interaction of G protein-coupled receptors (GPCRs) with β-arrestins (βarrs) is a critical mechanism that regulates the spatio-temporal pattern of receptor localization and downstream signaling. While the underlying mechanism governing GPCR-βarr interaction is primarily conserved and involves receptor activation and phosphorylation, there are several examples of receptor-specific fine-tuning of βarr-mediated functional outcomes. Considering the key contribution of conformational plasticity of βarrs in driving receptor-specific functional responses, it is important to develop and characterize novel sensors capable of reporting distinct βarr conformations in cellular context. Here, we design an intrabody version of a βarr-recognizing nanobody (nanobody32), referred to as intrabody32 (Ib32), in NanoLuc enzyme complementation assay format, and measure its ability to recognize βarr1 and 2 in live cells upon activation of a broad set of GPCRs. We discover that Ib32 robustly recognizes activated βarr1 and 2 in the plasma membrane as well as in the endosomes, and effectively mirrors βarr recruitment profile upon stimulation of GPCRs. We also design an Ib32 sensor for single-photon polarization microscopy with a change in linear dichroism as readout and demonstrate its utility for monitoring βarr activation upon stimulation of angiotensin receptor by its natural and biased agonists. Interestingly, when used side-by-side with a previously described sensor of βarr1 conformation known as Ib30, Ib32 uncovers distinct conformational signatures imparted on βarrs by different GPCRs, which is further corroborated using an orthogonal limited proteolysis assay. Taken together, our study presents Ib32 as a novel sensor to monitor βarr activation and leverages it to uncover conformational diversity encoded in the GPCR-βarr system with direct implications for improving the current understanding of GPCR signaling and regulatory paradigms.

## Introduction

G protein-coupled receptors (GPCRs), also referred to as seven transmembrane receptors (7TMRs), represent a large class of cell surface receptors and an important family of drug targets in the human genome^1–3^. Agonist-induced activation and phosphorylation of GPCRs leads to binding of β-arrestins (βarrs), which is a critical step in regulating receptor signaling and trafficking^4–7^. Interaction with GPCRs imparts an active conformation on βarrs that is manifested in the form of a significant inter-domain rotation between the N- and the C-domains, and the rearrangement of multiple loops in βarrs^8–14^. As the paradigm of agonist-induced βarr recruitment is typically conserved across nearly the entire repertoire of GPCRs, monitoring βarr activation may serve as a readout of receptor activation and ensuing downstream signaling. A number of biosensors of βarrs based on BRET and FRET have been designed and used to monitor activation dependent conformational changes in βarrs previously^15–20^. While some of these are capable of illuminating receptor-specific conformational signatures in βarrs, considering the ever-expanding layers of structural and functional complexities encoded in the GPCR-βarr system, additional biosensors amenable to cellular studies with easily accessible experimental set-up are still highly desirable.

Nanobodies have emerged as powerful tools in the recent years to probe novel aspects of GPCR activation and signaling, not only as conformational stabilizing chaperones for structural analysis but also as robust sensors for monitoring receptor activation^21–24^. In addition, several nanobodies targeting heterotrimeric G-proteins, originally described for structural investigation, have been successfully adopted to reveal novel aspects of spatio-temporal signaling and regulatory paradigms of GPCRs^24^. Still however, the use of intrabody sensors in the context of GPCR-βarrs is rather limited with only very few published examples^20,25,26^. Intrabody30 (Ib30), an intrabody derived from a synthetic antibody fragment (Fab30), has been developed and characterized to monitor agonist-induced GPCR-βarr1 interaction and trafficking in cellular context using confocal microscopy and NanoLuc enzyme-complementation-based assay^20,27^. Immunization of *Lama glama* with a pre-formed complex consisting of βarr1 and a chimeric β2-adrenergic receptor (β2V2R), and subsequent *in-vitro* screening, yielded several nanobodies that selectively recognize activated conformation of βarr1^28^. One of these, referred to as Nb32, was characterized in detail, first, in terms of promoting a fully-engaged β2V2R-βarr1 complex^28^, and then to visualize a β2V2R-G-protein-βarr1 endosomal signaling complex^29^. Considering the ability of Nb32 to selectively recognize activated βarr1, it represents a potential candidate to develop as an intrabody sensor of βarr activation in cellular context.

In this backdrop, here we describe a Nb32-based biosensor, referred to as intrabody32 (Ib32), in a NanoLuc/NanoBiT-based enzyme complementation format that is capable of reporting GPCR-induced activation of both isoforms of βarrs i.e., βarr1 and 2 for multiple GPCRs. When used in conjunction with a previously described Ib30 biosensor, Ib32 uncovers distinct conformations of βarrs induced by different GPCRs. We also design Ib32 sensors suitable for single-photon polarization microscopy with activation-dependent change in linear dichroism as readout and validate them on multiple GPCRs using balanced and biased-agonists. Collectively, Ib32-based sensors described here underscore the conformational diversity in GPCR-βarrs, and they may be useful tools for delineating previously unexplored complexities of GPCR signaling and regulation.

## Results

### Construct design and validation of Ib32 biosensor

Nb32 was identified to bind and stabilize β2V2R-βarr1 complex^28^, and subsequently used to determine the structure of β2V2R-G-protein-βarr1 endosomal signaling complex^29^. It binds to the N-domain of βarr1 in active conformation while it does not recognize the basal conformation of βarr1^28,29^ (Figure 1a), and therefore, it has the potential to be developed as a sensor of βarr activation. We retrieved the sequence of Nb32 from the previously determined structure and generated a set of constructs with N- and C-terminal fusion of the large and small fragments of the NanoLuc enzyme (i.e. LgBiT and SmBiT) following the principles of enzyme complementation assay (Figure 1b). These constructs are referred to as Ib32 (intrabody 32) as they are designed to express the Nb32 as a cytoplasmic protein. In order to validate and identify the optimal constructs, we performed enzyme complementation assay with each of these constructs in combination with the corresponding βarr1/2 versions described previously^27^ and the vasopressin receptor (V2R), a prototypical GPCR as a model system. We observed that a combination of LgBiT-Ib32 and SmBiT-βarr1 yielded maximal luminescence signal upon agonist-stimulation although other combinations also exhibited measurable signal (Figure 1c, Supplementary Figure 1a, b). A similar combination also worked with βarr2 although luminescence signal was relatively smaller than βarr1. In order to further corroborate these findings, we carried out the dose response curves of agonist-stimulation and observed a saturation response with increasing concentration of agonists with an EC_50_ value that is commensurate with that of βarr recruitment to V2R^20^ (Figure 1d, Supplementary Figure 1c, d). Furthermore, in order to directly visualize the Interaction of Ib32 with βarrs upon activation of V2R, we carried out co-immunoprecipitation experiments by pulling down either HA-Ib32 or Flag-V2R, and we observed a robust interaction of Ib32 with βarrs under agonist-stimulation conditions (Figure 1e, f and Supplementary Figure 2). Taken together, these data establish the ability of Ib32 to recognize activated conformation of βarrs in cellular context.

**Fig. 1.**
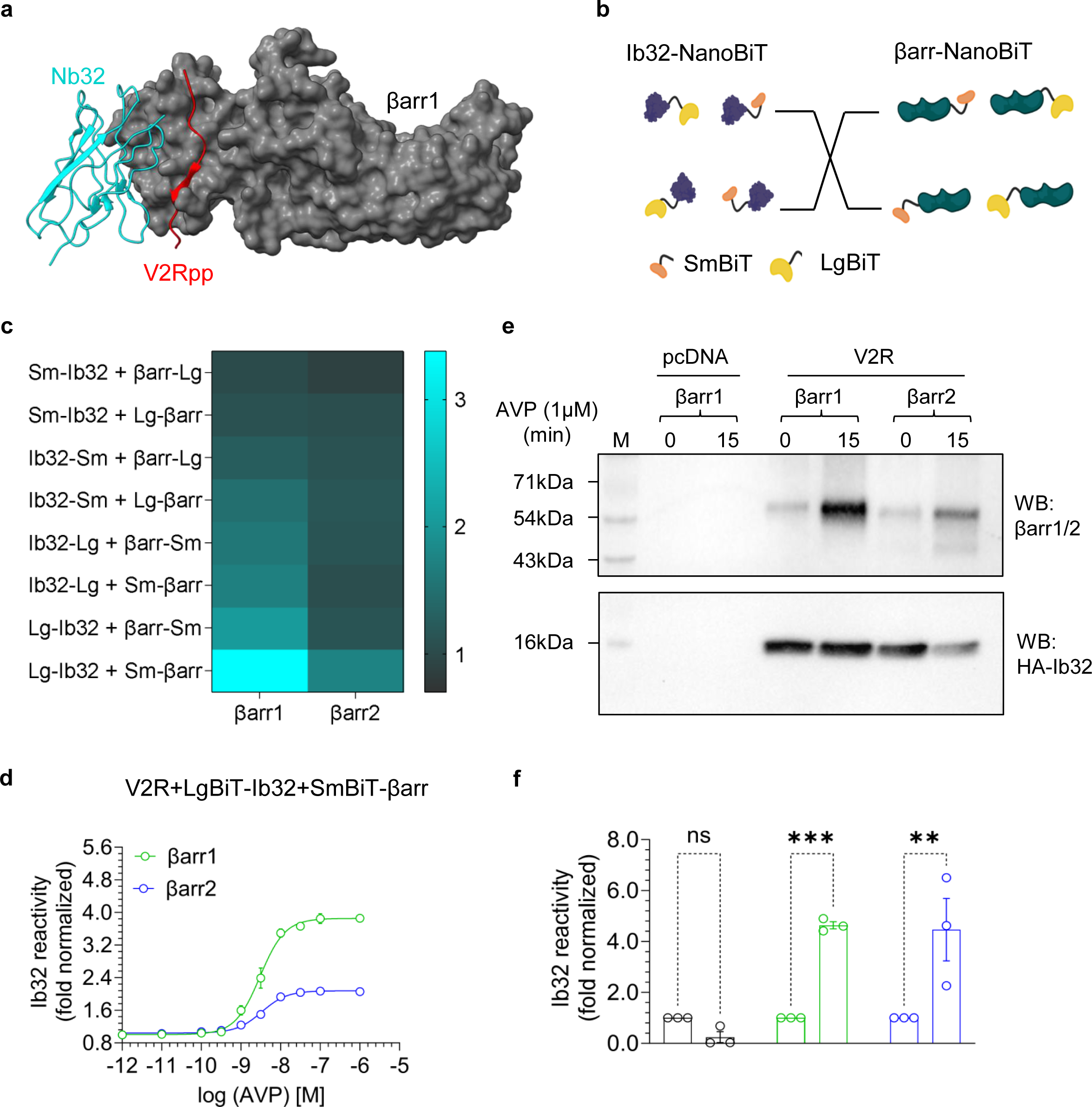
Construct design and validation of Ib32 as a sensor of βarr activation. **a,** Structural snapshot representing the binding interface of Nb32 on the N-domain of V2Rpp-bound βarr1, designed using ChimeraX based on the cryo-EM structure (PDB ID: 6NI2). **b,** Schematic depiction of the NanoBiT-based design of Ib32 and βarrs to identify the optimal constructs. **c,** Representation of the Ib32 reactivity data as a heat map for various combinations of the Ib32 and βarr1/2 NanoBiT constructs in response to stimulation of V2R with 1 µM AVP (mean±SEM; n=3; fold normalized with minimum concentration of each condition as 1). **d,** Dose response curve for Ib32 reactivity for βarr1/2 upon stimulation of V2R with increasing concentrations of AVP (mean±SEM; n=4; fold normalized with the minimum ligand concentration as 1). **e,** A representative blot showing Ib32 reactivity for βarr1/2 upon V2R stimulation as measured using co-immunoprecipitation assay. **f,** Densitometry analysis of the data presented in panel e, and the values represent mean±SEM of three independent experiment, normalized with respect to unstimulated condition treated as 1, analyzed using two-way ANOVA, Šídák’s multiple comparisons test.

### Ib32 as a sensor of βarr activation for multiple GPCRs

Next, we tested Ib32 sensor on a diverse set of GPCRs including complement and chemokine receptors, bradykinin B2 receptor, Angiotensin II type 1 receptor, muscarinic M2 receptor, kisspeptin receptor, motilin receptor, and the niacin receptor (GPR109A). We selected these receptors based on their relative propensities to interact with βarrs and binding modalities. For example, most of these receptors can be categorized as class A vs. B receptors based on stability of their interaction^30^ (class A: CXCR1, CXCR2, CXCR3, CXCR4, GPR109A; class B: C5aR1, AT1R, B2R), and they contain potential phosphorylation sites primarily in their carboxyl-terminus. We also used the muscarinic receptor subtype 2 (M2R) that interacts with βarrs primarily through the 3^rd^ intracellular loop instead of the carboxyl-terminus (M2R)^13^, and three β-arrestin-biased receptors namely the complement receptor C5aR2, decoy D6 receptor, and chemokine receptor CXCR7, which lack functional G-protein-coupling but robustly interact with βarrs^31,32^. While we observed a response for multiple GPCRs, the signal was most prominent for the B2R and AT1R (Figure 2a, Supplementary Figure 3a-d). We therefore carried out a dose response experiment for these two receptors and observed saturating responses with increasing agonist concentrations, and the EC_50_ values correspond well with βarr recruitment (Figure 2b, c, Supplementary Figure 3e-h). As different receptors exhibit a significant variation in their surface expression, it is plausible that a case-by-case optimization of the experimental conditions may allow a larger response even for those receptors, which do not appear to respond strongly in the screening panel presented in Figure 2a.

**Fig. 2.**
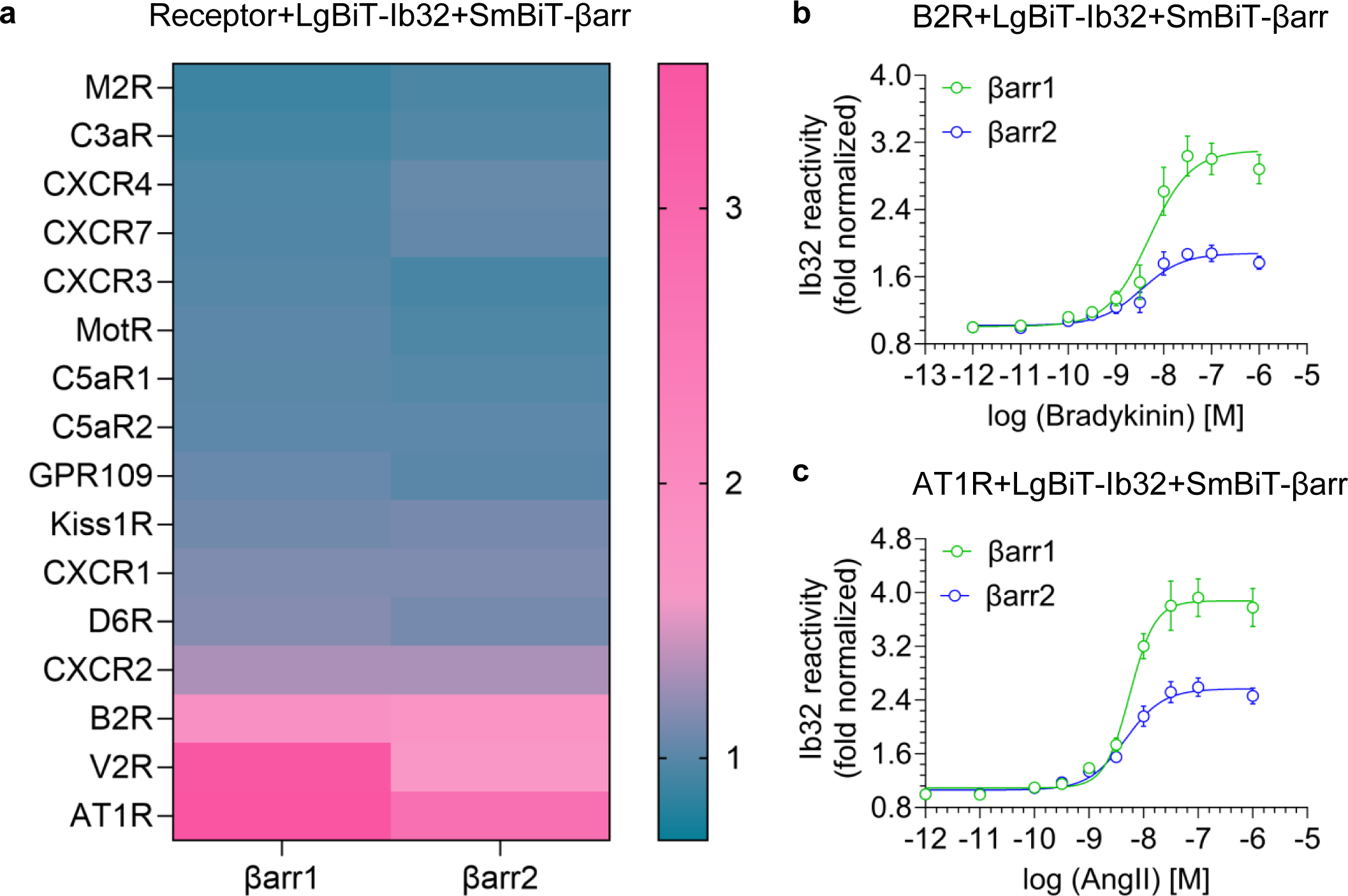
Ib32 as a sensor of βarr activation for multiple GPCRs. **a,** Representation of the Ib32 reactivity in the NanoBiT assay as a heat map for indicated receptors in response stimulation by their respective agonists. The values indicate average from 4-5 independent experiments, normalized with respect to unstimulated conditions for each receptors. **b-c,** Dose response experiment for Ib32 reactivity upon stimulation of B2R (panel **b**) and AT1R (panel **c**) with bradykinin and angiotensin II, respectively. Data represent mean±SEM from 3-4 independent experiments, normalized with respect to the response observed at minimum agonist concentration.

### Ib32 sensor for fluorescence-based linear dichroism microscopy

In order to broaden the application of Ib32 sensor for monitoring βarr activation in live cells, we designed a construct consisting of Ib32 with carboxyl-terminus fusion to a monomeric enhanced green fluorescent protein (meGFP) and a membrane-targeting isoprenylation signal peptide derived from H-Ras, referred to as Ib32-meGFP-H-Ras (Figure 3a). The design aims to exploit intrinsic optical anisotropy of fluorescent proteins^33^ in order to detect and characterize GPCR-βarr interactions. Briefly, fluorescent proteins anchored to the cell membrane often exhibit light absorption rates (and therefore fluorescence intensities) dependent on the direction of the linear polarization of the excitation light. This phenomenon, termed linear dichroism, can be used to determine the orientation of a fluorescent moiety with respect to the cell membrane^34^. As a result, a change in linear dichroism in a fluorescently-tagged molecule may provide an indication of a conformational change or protein-protein interaction^34^. We tested the Ib32-meGFP-H-Ras sensor on V2R and B2R using single-photon polarization microscopy and observed that stimulation resulted in a significant increase in linear dichroism for both βarr isoforms (Figure 3b-e). Similar to enzyme complementation assay, the Ib32-meGFP-H-Ras sensor exhibited a relatively larger change for βarr1 than βarr2, further corroborating the better ability of Ib32 to report βarr1 activation compared to βarr2.

**Fig. 3.**
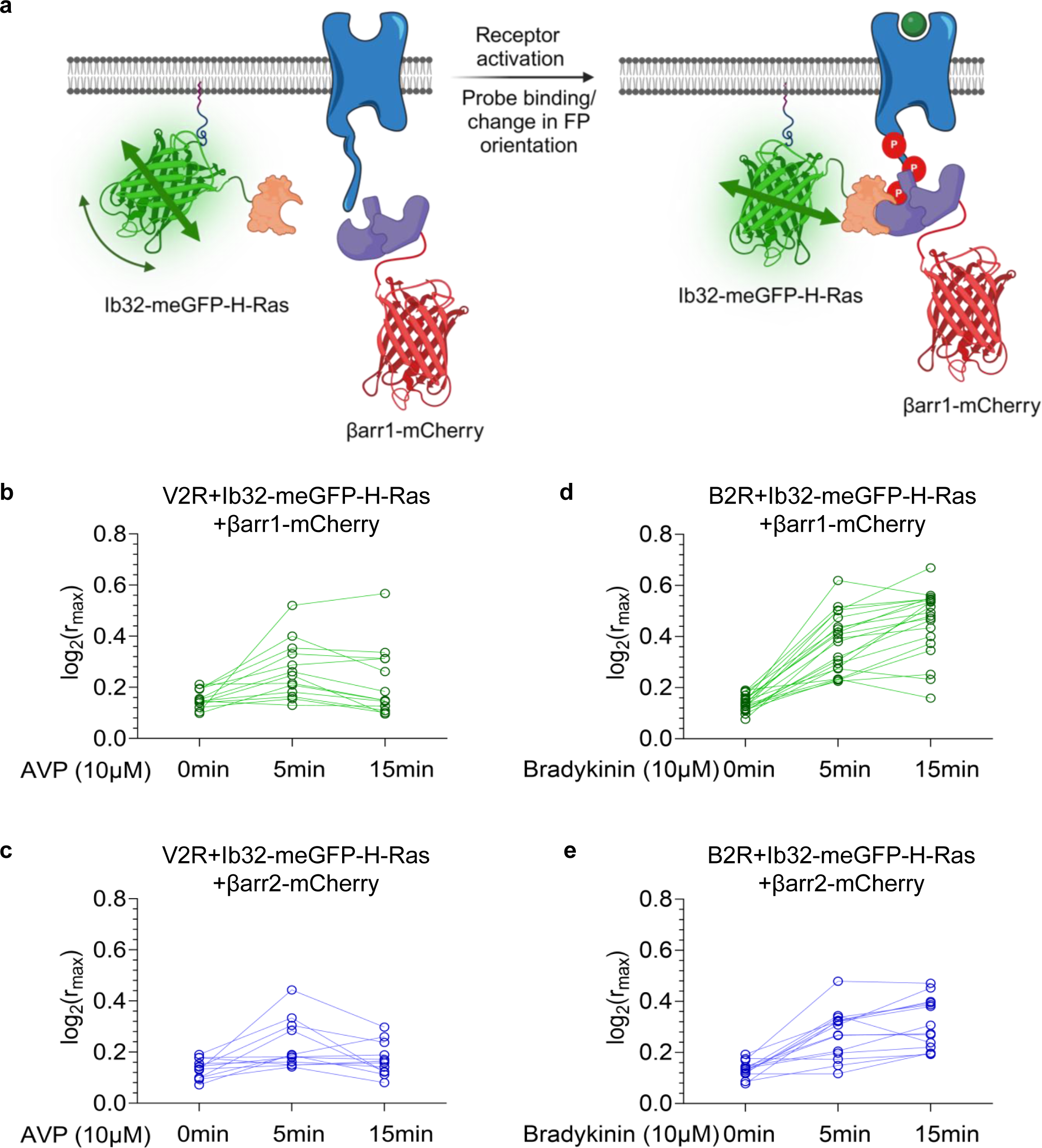
Ib32 sensor design and validation for fluorescence-based linear dichroism microscopy. **a,** Schematic of the Ib32 sensor as a fusion with meGFP and hRas designed to monitor a change in the linear dichroism as a readout of conformational change resulting from an orientational rearrangement of the meGFP moiety (prepared using BioRender). **b-e,** Agonist-induced changes in the linear dichroism of Ib32 sensor as measured upon stimulation of the V2R and B2R. The plots show the extent of linear dichroism {quantified as Log2(rmax)} upon binding of the Ib32-meGFP-H-Ras sensor to βarr1/2 in response to agonist-stimulation of V2R and B2R at indicated time-points. Each data point represents a single cell and means with 95% confidence intervals are indicated.

### Ib32 reactivity reports ligand pharmacology and compartmentalization of βarrs

Next, we tested the ability of Ib32 sensor to recognize βarr conformation upon stimulation of AT1R by a βarr-biased agonist, TRV027, vis-à-vis AngII, and we also measured βarr recruitment using a direct interaction assay based on the NanoBiT approach. We observed that TRV027 was relatively weaker in eliciting βarr recruitment compared to AngII (Figure 4a,c and Supplementary Figure 4a-d), and accordingly, Ib32 sensor mirrors the βarr recruitment pattern (Figure 4b,d). We also recapitulated an overall similar pattern for Ib32 sensor with respect to the change in linear dichroism as measured using single photon microscopy (Figure 4e-h). In the enzyme complementation assay, we likely observe a combination of βarr activation at the plasma membrane and endosomal compartment, and therefore, we next attempted to test if Ib32 preferentially recognizes βarrs at one of these locations. Here, we used either a plasma membrane-tethered LgBiT-CAAX or endosomal localized FYVE-SmBiT construct, and measured agonist-induced response. As presented in Figure 5a-d and Supplementary Figure 4e-l, we observed robust Ib32 reactivity in both cases; however, the fold response was higher in endosomal compartment than at the plasma membrane. However, it remains to be determined if the higher signal in the endosomal compartment reflects a distinct conformation of βarrs compared to the plasma membrane, or accumulation of internalized βarrs in the endosomes over time of experimental measurement. It is also interesting to note that in the case of V2R, Ib32 reactivity for βarr1 and 2 is similar at the plasma membrane but relatively stronger for βarr1 in the endosomal compartment, and future studies are warranted to explore this further.

**Fig. 4.**
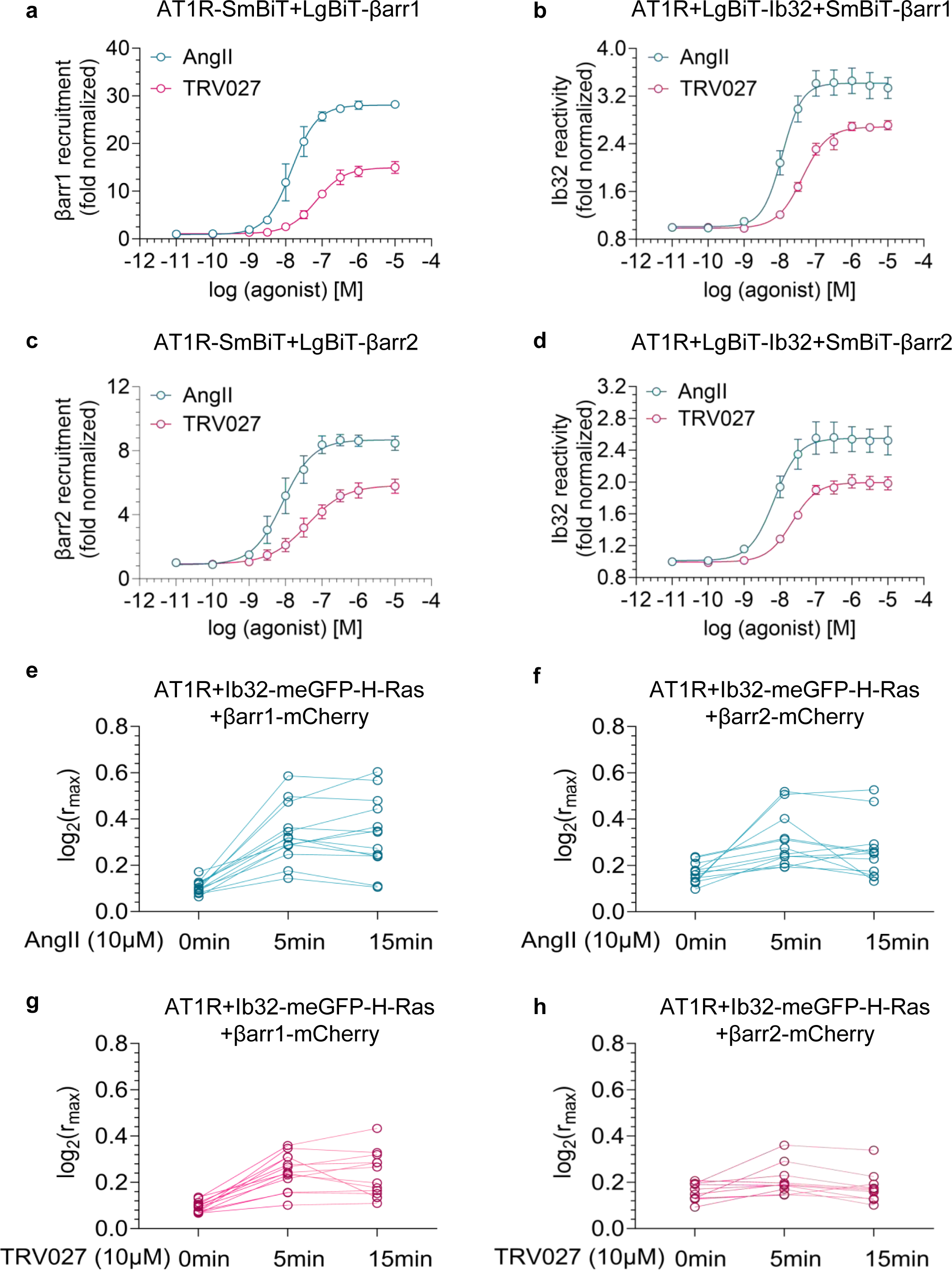
Ib32 sensor response aligns with ligand pharmacology. **a-d,** Dose response curves of βarr1/2 recruitment and Ib32 reactivity upon stimulation of AT1R with its endogenous agonist, angiotensin II (AngII) and a βarr-biased agonist, TRV027 (mean±SEM; n=3-4 independent experiments, normalized with respect to the response at minimum concentration of respective agonists treated as 1. **e-f,** Agonist-induced changes in the linear dichroism of Ib32 sensor upon stimulation of AT1R by angiotensin II and TRV027. The plots are showing the extent of linear dichroism {quantified as Log2(rmax)}, upon binding of Ib32-meGFP-hRas to βarr1/2 in response to agonist-stimulation for indicated time points. Each data point represents a single cell and mean with 95% confidence intervals are indicated.

**Fig. 5.**
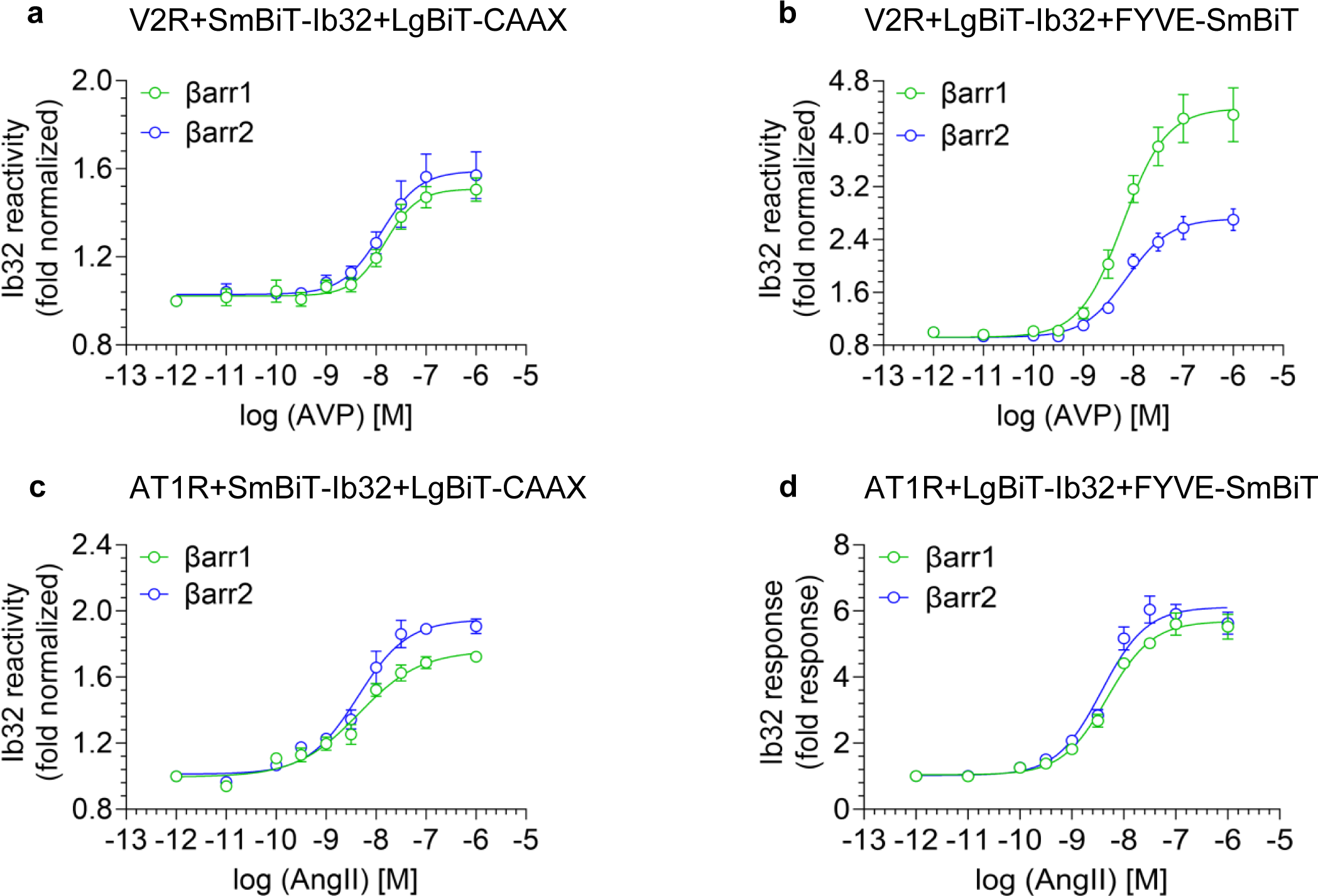
Ib32 reactivity at the plasma membrane and endosomes. **a-d,** Dose response curve for Ib32 reactivity at the plasma membrane using LgBiT-CAAX, and at the endosomes using FYVE-SmBiT constructs upon stimulation of V2R and AT1R with respective agonists. Data represent mean±SEM of 3-4 independent experiments, normalized with respect to the response observed at the minimum concentration of agonist.

### Ib32 reveals conformational diversity in GPCR-βarr complexes

We have previously developed and characterized an intrabody sensor referred to as intrabody30 (Ib30) based on an antibody fragment Fab30^8,20^, which also selectively recognizes GPCR-bound βarr1 conformation^15,20^. Thus, we compared the reactivity of Ib32 and Ib30 on a selected set of GPCRs to probe whether these two sensors recognize similar or different conformations of βarr1. We selected four distinct GPCRs namely V2R, the complement C5a receptor subtype 1 (C5aR1), Angiotensin II type 1 Receptor (AT1R), and the CXC chemokine receptor subtype 7 (CXCR7). Of these, Ib32 and Ib30 robustly recognized agonist-induced βarr1 conformation for the V2R while they both did not exhibit any measurable response for the CXCR7 (Figure 6a-d and supplementary Fig. 5a-c). The lack of response for CXCR7 is not due to absence of βarr1 recruitment as demonstrated using a NanoBiT assay reporting agonist-induced βarr1 recruitment to the receptor under similar experimental conditions (Figure 6a). These data suggest that upon binding to CXCR7, βarr1 adopts a conformation that is significantly different from that induced by V2R. Interestingly, Ib32 robustly recognizes βarr1 upon stimulation of AT1R but fails to recognize βarr1 for C5aR1 (Figure 6a-d). On the other hand, Ib30 displays a pattern that is nearly-reverse of Ib32 (Figure 6a-d). Expectedly, AT1R and C5aR1 display robust βarr1 recruitment under similar experimental conditions. Taken together, these data suggest at least four distinct conformations of βarr1 upon its interaction with V2R, C5aR1, AT1R and CXCR7, and therefore, underscores the conformational diversity displayed by βarrs upon their interaction with GPCRs.

**Fig. 6.**
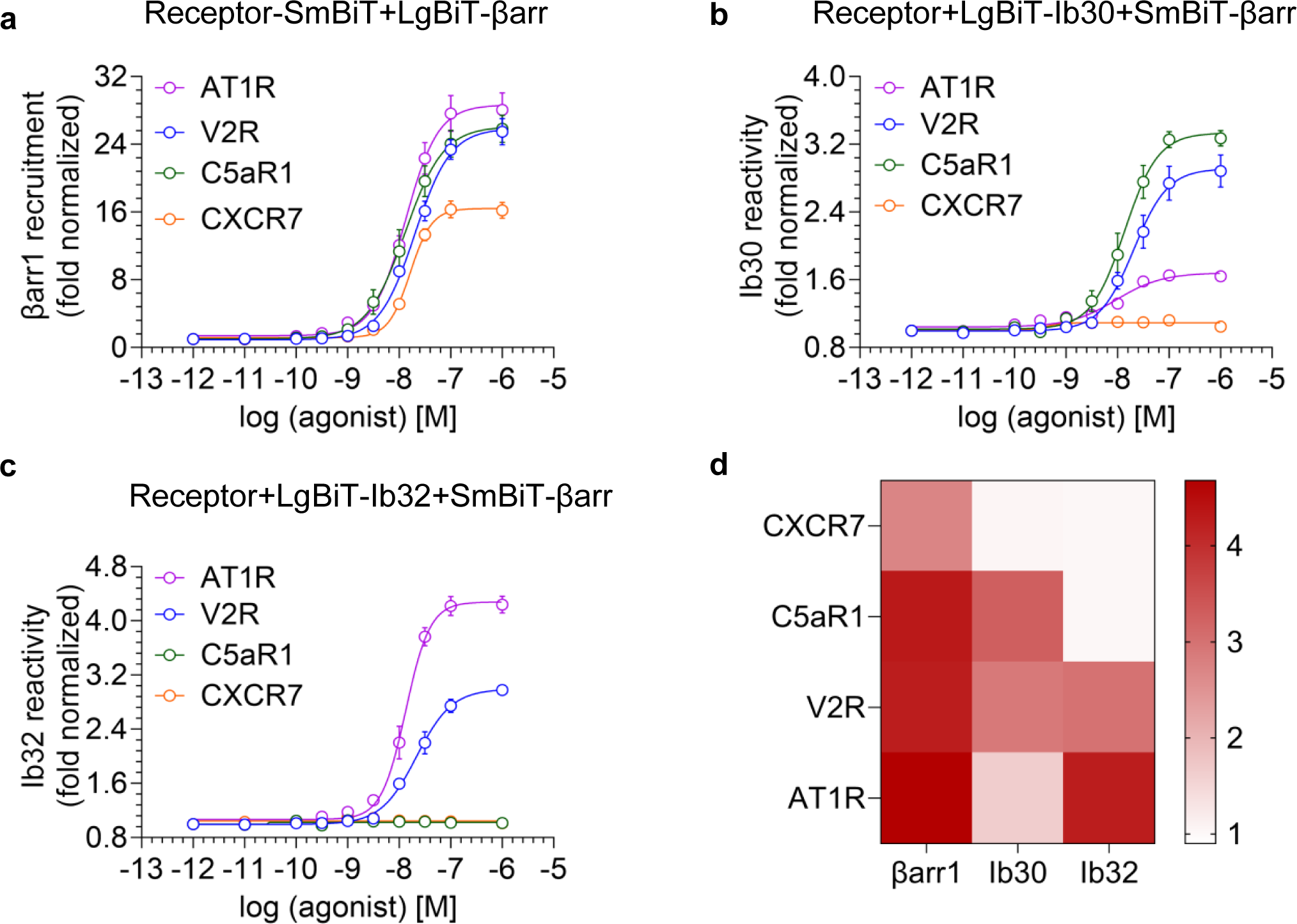
Ib32 sensor reveals conformation diversity in GPCR-βarr complexes. **a-c,** βarr1 recruitment, Ib30 reactivity, and Ib32 reactivity depicted in panel a, b, and c, respectively, for indicated receptors upon stimulation with increasing concentrations of the corresponding agonists. Data represent mean±SEM of 3-4 independent experiments, normalized with respect to the minimum concentration of corresponding agonists. **d,** Representation of βarr1 recruitment, Ib30 and Ib32 reactivity responses observed at maximal agonist concentrations for the indicated receptors as a heat map.

### Phosphorylation sites in the B2R driving Ib32 reactivity

As Ib32 exhibits strongest signal in the case of B2R, we next set out to identify the contribution of distinct phosphorylation sites in the B2R driving βarr interaction and conformation activation as recognized by Ib32. We generated a series of phosphorylation site mutants of B2R as depicted in Figure 7a, and measured their ability to recruit βarrs upon agonist-stimulation and the reactivity of Ib32 using the NanoBiT assay in parallel. We observed that most of the mutants maintained βarr recruitment and Ib32 reactivity to the levels comparable to wild-type receptor at saturating dose of agonist (Figure 7b-c and supplementary Figure. 5d-g). However, a combination of Thr^372^Ala and Ser^373^Ala nearly abolished Ib32 reactivity despite maintaining robust βarr recruitment (Figure 6d-g and Supplementary Figure 5h-k). These data suggest that phosphorylation of Thr^372^ and Ser^373^ are critical for imparting a βarr conformation that is recognized by Ib32.

**Fig. 7.**
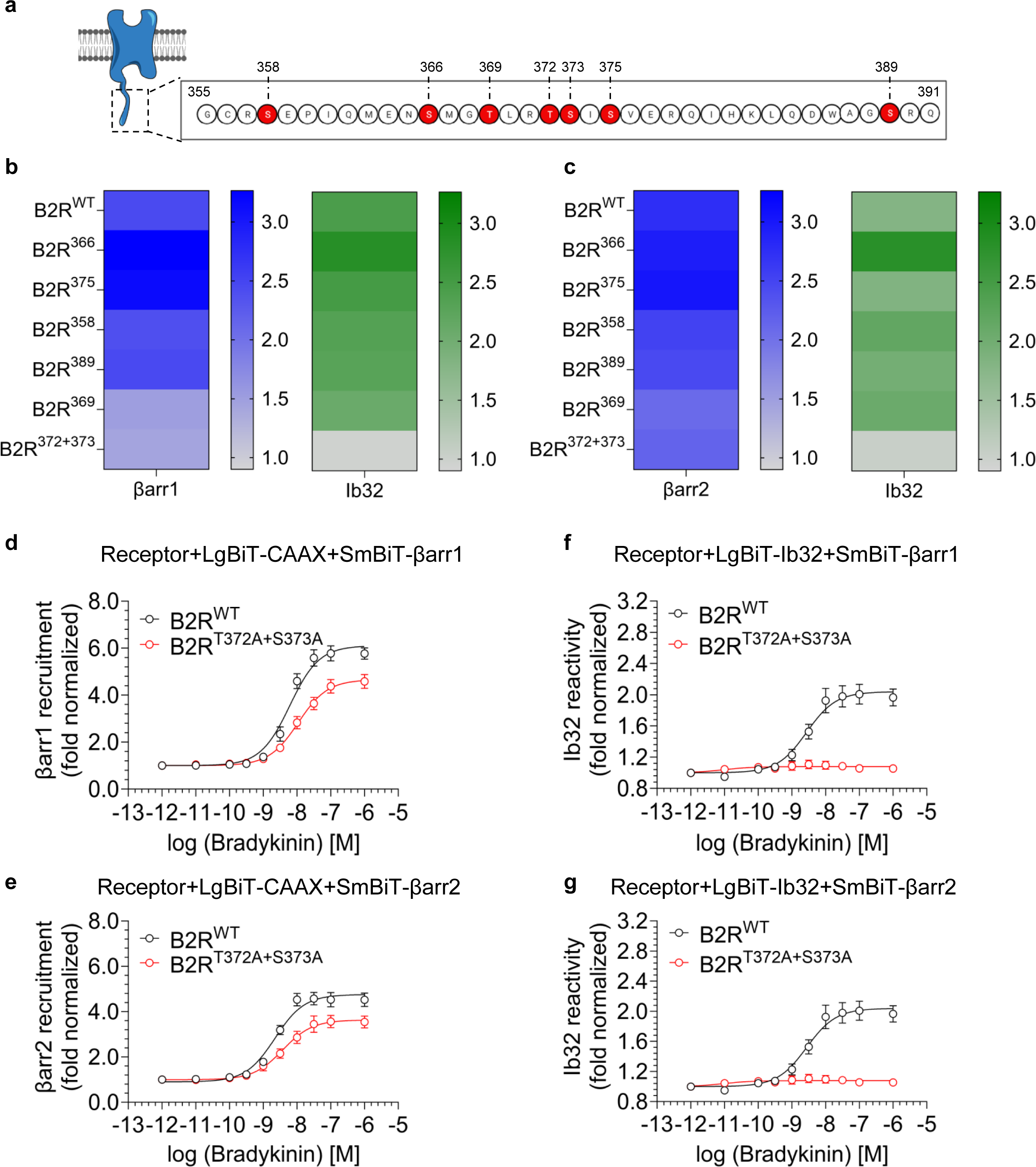
Contribution of different phosphorylation sites in B2R on Ib32 sensor response. **a,** Schematic representation showing the potential phosphorylation sites in the carboxyl-terminus of B2R (prepared using BioRender). **b, c,** Heatmap representation of βarr1/2 recruitment and Ib32 reactivity for the phosphorylation site mutants of B2R at saturating agonist concentration (1μM). The values indicate average of three independent experiments, normalized with respect to the unstimulated condition. **d-g,** Dose response experiment for βarr1/2 recruitment (panel d and e) and Ib32 reactivity for βarr1 (panel f) and βarr2 (panel g) with B2R_WT_ and B2R^T372A+S373A^. Data represent mean±SEM of 3-4 independent experiments, normalized with respect to the minimum concentration of the agonist.

### Limited proteolysis corroborates conformation diversity in βarr activation

Finally, we employed a limited proteolysis assay to corroborate the conformational diversity imparted by different GPCRs on βarrs. For this, we used phosphopeptides derived from the carboxyl-terminus of multiple GPCRs to activate βarrs *in-vitro* (Supplementary Figure. 6). In these experiments, we first activated purified βarr1 using saturating concentrations of phosphopeptides followed by limited trypsin proteolysis and monitored the kinetics of proteolysis as a readout of βarr activation. We observed that the kinetics of limited proteolysis for V2Rpp vs. B2Rpp were different from each other wherein B2Rpp binding shows a slower rate of proteolysis compared to that of V2Rpp (Figure 8a,c). Moreover, phosphopeptides derived from different GPCRs and containing distinct phosphorylation patterns also exhibited differential kinetics of limited proteolysis (Figure 8b,d). Taken together with the Ib32 reactivity data, these findings underscore the conformational fine-tuning in βarr1 upon activation by distinct phosphorylation patterns harboured in different GPCRs.

**Fig. 8.**
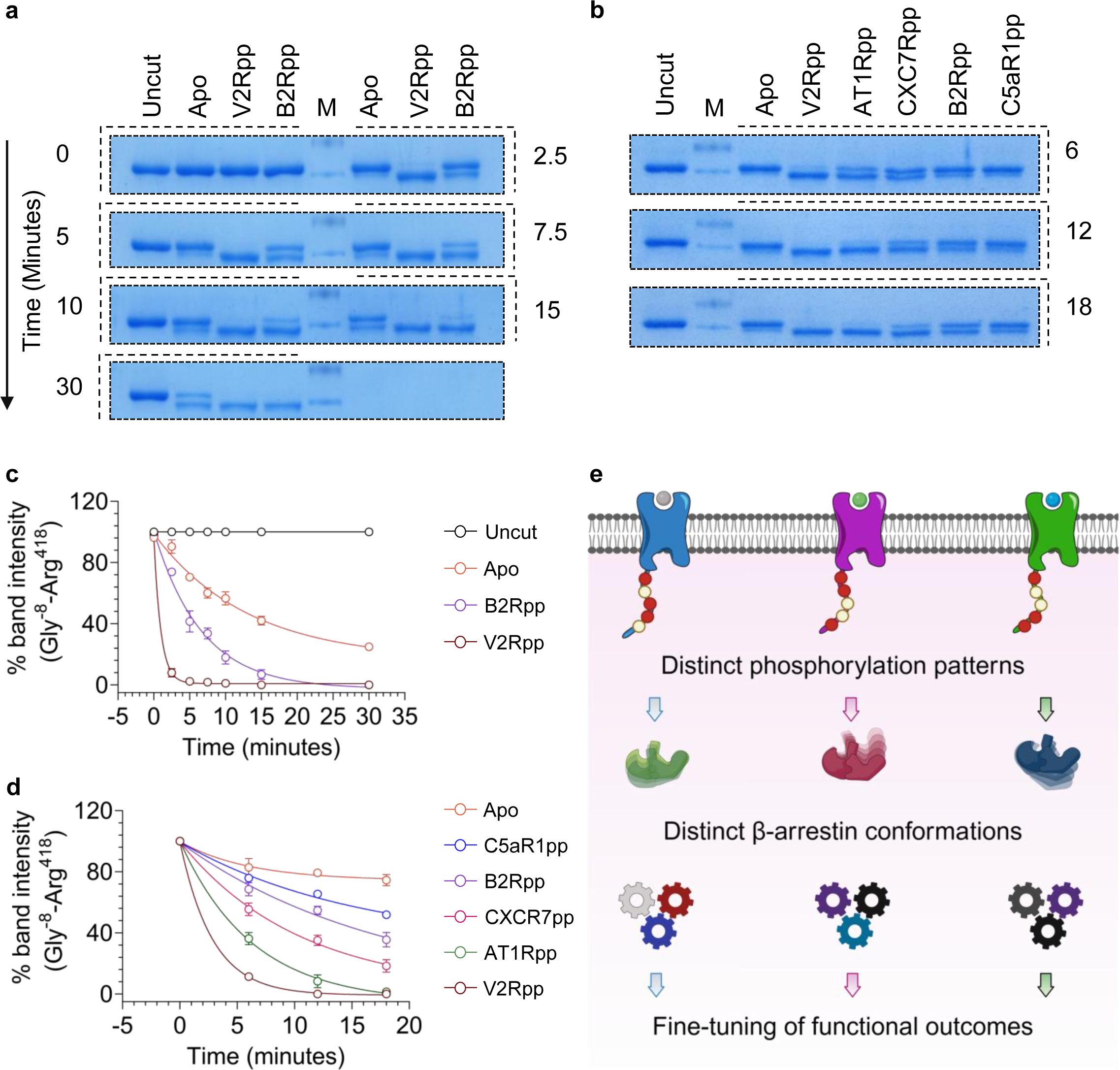
Limited proteolysis of GPCR phosphopeptide-activated βarr1. **a-b,** Representative images for limited trypsin proteolysis assay carried out on purified βarr1 after their activation with saturating concentrations (ten-fold molar excess) of the indicated phosphopeptides. The proteolytic fragments were separated by SDS-PAGE and visualized using coomassie brilliant blue staining. **c-d,** Densitometry-based quantification of the band intensities corresponding to the full-length βarr1. Data represent mean±SEM of four independent experiments, normalized as % with respect to the starting intensity (i.e. undigested). **e,** A schematic representation depicting diverse conformational signatures imparted on βarrs upon their interaction and activation by different GPCRs (prepared using BioRender).

## Discussion

In this study, we develop Ib32 as a novel sensor to monitor βarr activation upon their interaction with GPCRs, and employ it in cellular context to demonstrate conformational diversity in GPCR-βarr complexes. It is important to note that Ib32 is not universal for the entire spectrum of GPCRs and its utility for different receptors should be evaluated individually. Interestingly, for some receptors such as CXCR2, Ib32 yields clear response upon agonist-stimulation although the window of detection is relatively weaker compared to other GPCRs. This may result from a lower expression level of these receptors, affinity and/or conformation of βarr interaction arising from their phosphorylation pattern, and it warrants additional studies in future. It is interesting to note that Ib32 sensor reports βarr1 activation with a stronger signal compared to βarr2 for some receptors while it is equally efficient for others. This is in line with previous studies based on intramolecular FlAsH-BRET sensors of βarrs, which have illuminated receptor-specific βarr conformational signature^16,17^, and also conformational differences between the βarr isoforms when activated by the same receptor^15,35,36^.

We observed that Ib32 sensor displays a robust change in linear dichroism upon activation of βarrs by three different GPCRs tested here. Although the direction of changes in linear dichroism was similar for all three receptors and βarr isoforms, we observed a slightly better signal for βarr1 vs. βarr2. Moreover, the linear dichroism changes were similar upon stimulation of AT1R by AngII and TRV027 with the signal for the latter being smaller, which corresponds with their βarr recruitment profile. A number of previous studies have demonstrated that biased agonists typically induce a distinct βarr conformation compared to unbiased or balanced agonists^37^, and therefore, it is likely that Ib32 sensor, either in the NanoBiT format or in linear dichroism setting, is not able to differentiate between such conformational differences. A series of recent studies have proposed different phosphorylation signatures and motifs in GPCRs as critical determinants of βarr interaction and activation^12,14,38–40^. Moreover, several biochemical and cellular studies using site-directed mutagenesis and functional assays, have also linked specific βarr conformations to downstream responses^27,28,41–44^. The data presented here with Ib32 sensor underscores additional level of conformation diversity in βarrs that is imparted and fine-tuned in receptor-specific manner (Figure 8e), and future studies focused on direct structural visualization of these complexes may provide further insights.

In conclusion, we present the development and characterization of Ib32 as a genetically-encoded sensor of βarr recruitment and activation for selected GPCRs. Taken together with a previously described Ib30 sensor, it demonstrates the existence of distinct conformational signatures in βarrs for different GPCRs, and going forward, it should complement the existing biosensors for visualizing novel aspects of GPCR-βarr interaction and activation.

## Acknowledgements

This work is supported by an Indo-Czech collaborative research grant from the Department of Science and technology, Government of India (File No. DST/INT/Czech/P-03/2019) and Science and Engineering Research Board (SERB) (SPR/2020/000408). In addition, the research program in the laboratory of A.K.S. is supported by the Senior Fellowship of DBT Wellcome Trust India Alliance (IA/S/20/1/504916) and Indian Council for Medical Research (F.NO.52/15/2020/BIO/BMS). A.K.S. is a recipient of the Sonu Agrawal Memorial Chair from IIT Kanpur. The research on this project in J.L.’s laboratory is supported by an InterExcellence/Inter-Action grant LTAIN19167 by the Ministry of Education, Science and Sports of the Czech Republic. We thank Pragyashree Bhowmick for assisting with limiting proteolysis experiments and Manisankar Ganguly for preparing Figure 1a.

## Authors’ contribution

PS and VM carried out Ib32 construct design, sub-cloning, and preliminary characterization; PS performed the cellular assays with help from AD, NZ and SM; PM prepared the Nb32-meGFP-hRas and IB30-meGFP-hRas constructs; MKY carried out the limited proteolysis assay with help from GM and NR; VM carried out the polarization microscopy experiments and analyzed the data under the supervision of JL; all authors contributed to data interpretation and manuscript writing; JL and AKS supervised the overall project.

## Materials and methods

### General reagents, plasmids, and cell culture

Most of the standard reagents were purchased from Sigma Aldrich unless specified otherwise. Dulbecco′s Modified Eagle′s Medium (DMEM), Phosphate Buffer Saline (PBS), Trypsin-EDTA, Fetal-Bovine Serum (FBS), Hank’s Balanced Salt Solution (HBSS), and Penicillin-Streptomycin solution were purchased from Thermo Fisher Scientific. HEK-293 cells were purchased from ATCC and maintained in 10% (v/v) FBS (Gibco, Cat. no. 10270-106) and 100 U ml^-1^ penicillin and 100 µg ml^-1^ streptomycin (Gibco, Cat. no. 15140122) supplemented DMEM (Gibco, Cat. no. 12800-017) at 37 °C under 5% CO_2_. The expression constructs for the receptors, βarrs, NanoBiT-CAXX, and Ib30 have been described previously^13,20,27,31,32,45–48^. Ib32 constructs in NanoBiT format as described in Figure 1b were generated by sub-cloning the Ib32 coding region in pCAGGS vector as described previously for Ib30^27^. For linear dichroism imaging, the Ib32-mEGFP-hRas construct was cloned into mEGFP-hRas cassette in a pcDNA3.1(+) vector. B2R phosphorylation site mutants mentioned in the manuscript were generated by site-directed mutagenesis using Q5 Site-directed mutagenesis kit (NEB, Cat. No. E0554). All DNA constructs used in this study were verified by sequencing from Macrogen. Arginine Vasopressin (AVP), Angiotensin (AngII), Bradykinin, Motilin-22, and Kisspeptin-10 were synthesized from GenScript. Niacin and Carbachol were purchased from Himedia (Cat. no. TC157) and Cayman Lifesciences respectively (Cat. no. 14486). C5a, C3a, CCL7, CXCL8, CXCL11, and CXCL12 were purified from the *E. coli* BL21(DE3) cells following the protocols described previously^49–51^.

### Receptor surface expression

Surface expression of the receptors in various assays was measured using a whole cell-based surface ELISA protocol reported previously^52^. Briefly, transfected cells were seeded in 0.01% poly-D-Lysine pre-treated 24-well plate at a density of 2×10^5^ cells well^-1^. Post 24 h of incubation, once cells were washed with ice-cold 1X TBS, followed by fixation with 4% PFA (w/v in 1X TBS) on ice for 20 min. Fixed cells were then washed with 1X TBS for three times followed by blocking with 1% BSA (w/v in 1X TBS) at room temperature for 90 min. Post blocking, 200 μl anti-FLAG M2-HRP was added and incubated for 90 min (prepared in 1% BSA, 1:10,000) (Sigma, Cat. no. A8592). Post blocking, cells were washed with 1% BSA (prepared in 1X TBS) three times to remove unbound antibodies. Signal was developed by treating cells with 200 μl TMB-ELISA (Thermo Scientific, Cat no. 34028) until the light blue colour appeared, and the reaction was quenched by transferring the solution to a 96-well plate containing 100 μl 1 M H_2_SO_4_. The absorbance of the signal was measured at 450 nm using a multi-mode plate reader. After taking absorbance of the signal for FLAG tagged receptor, cells were washed once with 1X TBS followed by incubation with 0.2% Janus Green (Sigma; Cat. no. 201677) w/v for 15 min. Afterwards, Janus Green was aspirated followed by washing with distilled water to remove the excess stain. To elute the stain, 800 μl of 0.5 N HCl was added. To record the absorbance of Janus, 200 μl of the eluate was transferred to a 96-well plate and absorbance was measured at absorbance 595 nm. Data were analyzed by calculating the ratio of absorbance at 450/595 followed by normalizing the value of pcDNA-transfected cells reading as 1, and the values were plotted using GraphPad Prism v 9.5.0 software.

### NanoBiT enzyme complementation assay

NanoBiT assay to monitor GPCR-βarr interaction and Ib32/Ib30 reactivity was carried out following the previously described protocols^13,20,27,31,45–47,51^. For the Ib32 reactivity assay described here for the first time, various combinations of LgBiT and SmBiT tagged at the N-terminus and C-terminus of ꞵarr1/2 and Ib32 were screened to obtain the optimal combination. HEK-293 cells were transfected with 3 µg of receptor, 2 µg of βarr1/2 (with indicated fusion of LgBiT and SmBiT) and 5 µg of Ib32 (with indicated fusion of LgBiT and SmBiT) using the transfection reagent polyethyleneimine (PEI) linear at DNA: PEI ratio of 1: 3. After 16-18 h of transfection, cells were harvested and resuspended in the NanoBiT assay buffer containing 1X HBSS, 0.01% BSA, 5 mM HEPES pH 7.4, and 10 μM coelenterazine (GoldBio, Cat. no. CZ05). Afterwards, resuspended cells were seeded at a density of 1×10^5^ cells well^-1^ in an opaque flat bottom 96-well plate. Post-incubation, basal level luminescence readings were recorded, followed by addition of 1 μM arginine vasopressin (AVP). Luminescence upon stimulation was recorded for 20 cycles using a multi-mode plate reader (BMG Labtech). We observed the maximal response with N-terminally LgBiT tagged Ib32 and βarr with SmBiT at the N-terminus, and this combination was used in subsequent experiments. A similar combination of SmBiT-tagged βarr and LgBiT fused Ib30 was also used to probe Ib30 reactivity for βarr activated by the indicated receptors. For analysis, stimulated readings were normalized with respect to the signal of minimal ligand concentration as 1 and plotted using nonlinear regression analysis in GraphPad Prism v 9.5.0 software. For βarr recruitment assay, HEK-293 cells were transfected with 3.5 μg of receptor fused with SmBiT at the C-terminus and 3.5 μg of N-terminally LgBiT tagged βarr1/2 followed by the same procedure as described above. In order to measure βarr recruitment to B2R phosphorylation site mutants, bystander NanoBiT-based was used where HEK-293 cells were transfected with 5 μg of receptor, 2 μg of N-terminally SmBiT fused βarr1/2, and 5 μg LgBiT-CAAX construct. In the experiments measuring plasma membrane and endosomal recruitment of Ib32, cells were transfected with 2µg of the receptor and 2 µg of βarr1 together with either 4 µg of SmBiT-Ib32 + 4 µg of LgBiT-CAAX, or 4 µg of LgBiT-Ib32+4 µg of FYVE-SmBiT.

### Co-immunoprecipitation assay

In order to measure the Ib32 reactivity to βarr1/2 upon stimulation of V2R, a co-immunoprecipitation (co-IP) assay was carried out following a previously described protocol^53^. Briefly, HEK-293 cells were co-transfected with 3 μg of N-terminally FLAG tagged V2R, 1.5 μg of βarr1 or 2 μg of βarr2 and 4 μg of Ib32 fused with HA tag at the C-terminus. Post 48 h of transfection, complete DMEM is replaced with incomplete DMEM for serum starvation for 6 h. Afterwards, cells were stimulated with 1 μM AVP for 15 min, harvested, resuspended in 100 μl lysis buffer (20 mM HEPES pH 7.4, 450 mM NaCl, 0.1 mM PMSF, 0.2 mM Benzamidine, and 1X Phosphatase inhibitor cocktail), and lysed by dounce homogenization. Post-lysis, sample was solubilized with 1% L-MNG (maltose neopentyl glycol) for 1 h at room-temperature. For the experiment presented Fig. 1e, supernatant was allowed to bind with the anti-HA antibody bound beads, followed by washing with wash buffer (20mM HEPES, pH 7.4, 100mM NaCl) and elution using 30μl 2X SDS reducing dye. For the co-IP data presented in Supplementary Figure. 1e, M1-FLAG beads were used, and the beads were subjected to alternative washes with low salt buffer (20 mM HEPES pH 7.4, 150 mM NaCl, 2 mM CaCl_2_, 0.01% L-MNG) and high-salt buffer (20 mM HEPES pH 7.4, 350 mM NaCl, 2 mM CaCl_2_, 0.01% L-MNG), followed by elution with FLAG-EDTA buffer (20 mM HEPES pH 7.4, 150 mM NaCl, 2 mM EDTA, 0.01% MNG, 250 μg ml^-1^ FLAG peptide). Samples were then subjected to separation by SDS-polyacrylamide gel electrophoresis and transferred to PVDF (Polyvinylidene fluoride) membrane. Afterwards, βarr1/2, Ib32, and the receptors were probed using βarr1/2 antibody (1:5000, CST, Cat. no. 4674), monoclonal anti-rabbit IgG peroxidase coupled antibody, anti-HA antibody (dilution-1:5000; Santa-Cruz; cat. no. sc-805), and anti-FLAG peroxidase coupled antibody (1:5000, Sigma-Aldrich, Cat. no. A8592), respectively. Signals were detected using a Chemiluminescence Documentation imaging system (Bio-Rad), densitometry-based quantification was carried out using ImageJ software suite, data were plotted using GraphPad Prism v 9.5.0 Software.

### Ib32 sensor design and experimental details for single photon microscopy

The Ib32-meGFP-H-Ras construct was prepared by PCR amplification of the Ib32 gene, restriction digestion by XbaI/XhoI, and cloning into the corresponding sites of an meGFP-H-Ras cassette in a pcDNA3.1(+) vector, prepared by gene synthesis (GenScript). Prior to polarization microscopy imaging, cells (HEK-293) were plated in 8-well µ-slides (iBidi, Germany) and transfected using Lipofectamine 3000 (Thermo Fisher Scientific) and a procedure recommended by the manufacturer. Transfection was carried out using 500 ng each of plasmids encoding Ib32-meGFP-H-Ras, the studied GPCR (V2R, B2R or AT1R), and βarr1-mCherry or βarr2-mCherry. After transfection, cells were incubated at 37 °C overnight. For activations, agonists were manually added by pipetting, to a final concentration of 10 µM. Cells were observed by single-photon polarization microscopy prior to addition of an agonist, and 5 and 15 min after adding an agonist.

Single-photon polarization microscopy was performed as described previously^34,54^ using an Olympus FV1200 confocal microscope equipped with a polarization modulator (RPM-2P, Innovative Bioimaging) alternating the direction of the excitation light polarization between acquisition of subsequent pixels. Ib32-meGFP-H-Ras was observed using laser light of 488 nm wavelength and 80 µW intensity, through a 40X water immersion objective lens (UApoN340, NA1.15, Olympus, Japan). Images containing information on fluorescence intensities observed with both horizontal and vertical excitation polarizations were deconvolved and quantitatively processed in ImageJ/Fiji, using publicly available macros described previously^54^. The extent of linear dichroism was characterized by a value of the maximum dichroic log ratio {log_2_(*r*_max_)}, defined as the base-2 logarithm of the ratio of fluorescence intensities observed with light polarized horizontally and vertically in the image, in a section of the cell membrane oriented horizontally in the image. For each combination of constructs and experimental conditions, at least 10 cells were observed and analyzed. The resulting values of log_2_(*r*_max_) were statistically analyzed and plotted using GraphPad Prism v 9.5.0 software.

### Limited trypsin proteolysis assay

As an orthogonal approach to measure activation-dependent βarr conformational change, we employed a limited trypsin proteolysis assay as described previously^47,55,56^ with minor modifications. Briefly, 5 µg of βarr1 was incubated with 10 molar excess of either V2Rpp or B2Rpp for 40 min in ice-cold reaction buffer (20 mM HEPES pH 7.4, 100 mM NaCl). Following activation, 20 µL aliquot was collected as the time zero control and TPCK-treated trypsin was added at a 1: 200 (w/w) trypsin: βarr1 ratio, and the mixture was incubated at 37 °C. 20 µl sample was withdrawn at different time intervals and quenched with 5 µl SDS-protein loading dye. 10 µl sample was run on a 12% SDS-PAGE for quantitative analysis. The decrease in intensity of the Gly^-8^ to Arg^418^ (48kDa) band was quantified by densitometry and data were plotted using GraphPad Prism v 9.5.0 software.

