## Supplemental Figures for "A genetically-encoded nanobody sensor reveals conformational diversity in β-arrestins orchestrated by distinct seven transmembrane receptors"

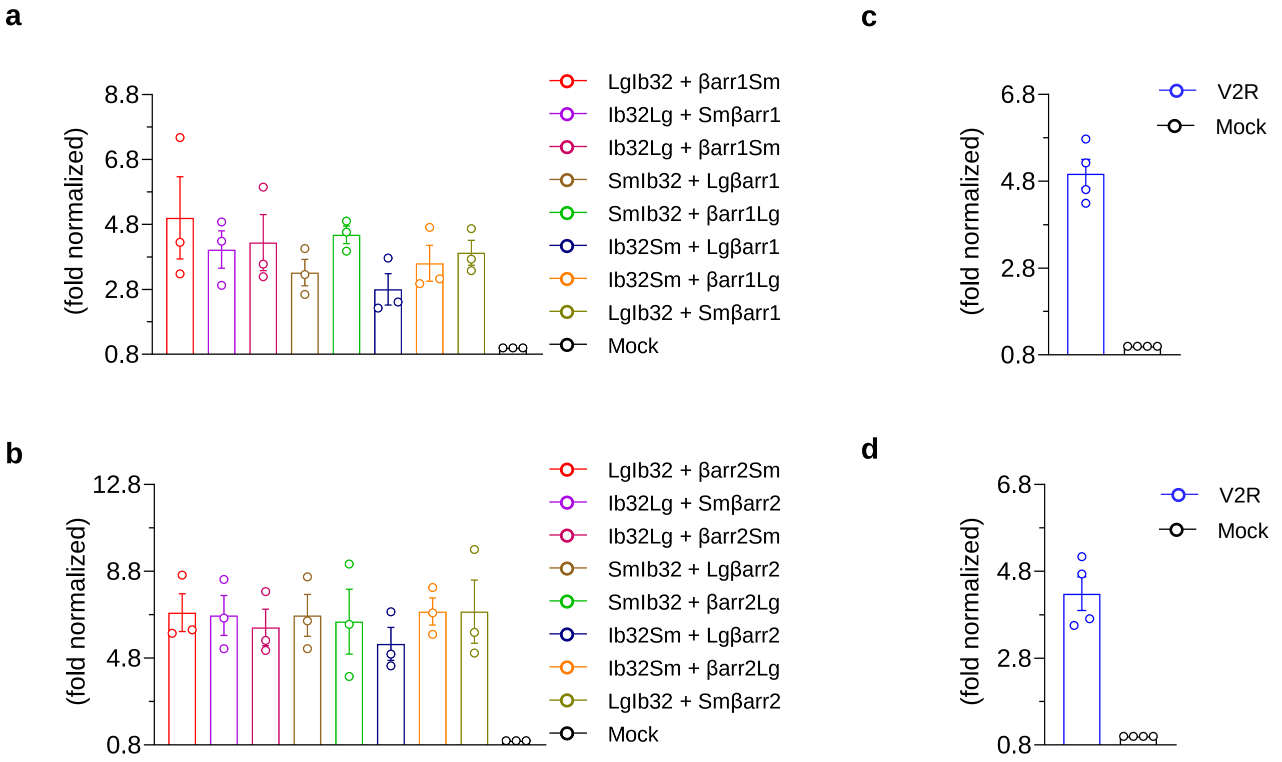


**Supplementary Figure. 1. Surface expression of V2R. a, b,** Surface expression of V2R in the Ib32 reactivity assay data presented in Figure 1c, measured using whole cell ELISA. Data represent mean±SEM of three independent experiments, normalized as fold over mock-transfected condition. **c, d,** Surface expression of V2R in the Ib32 reactivity dose response experiment presented in Figure 1d, measured using whole cell ELISA. Data represent mean±SEM of four independent experiments, normalized as fold over mock-transfected condition.

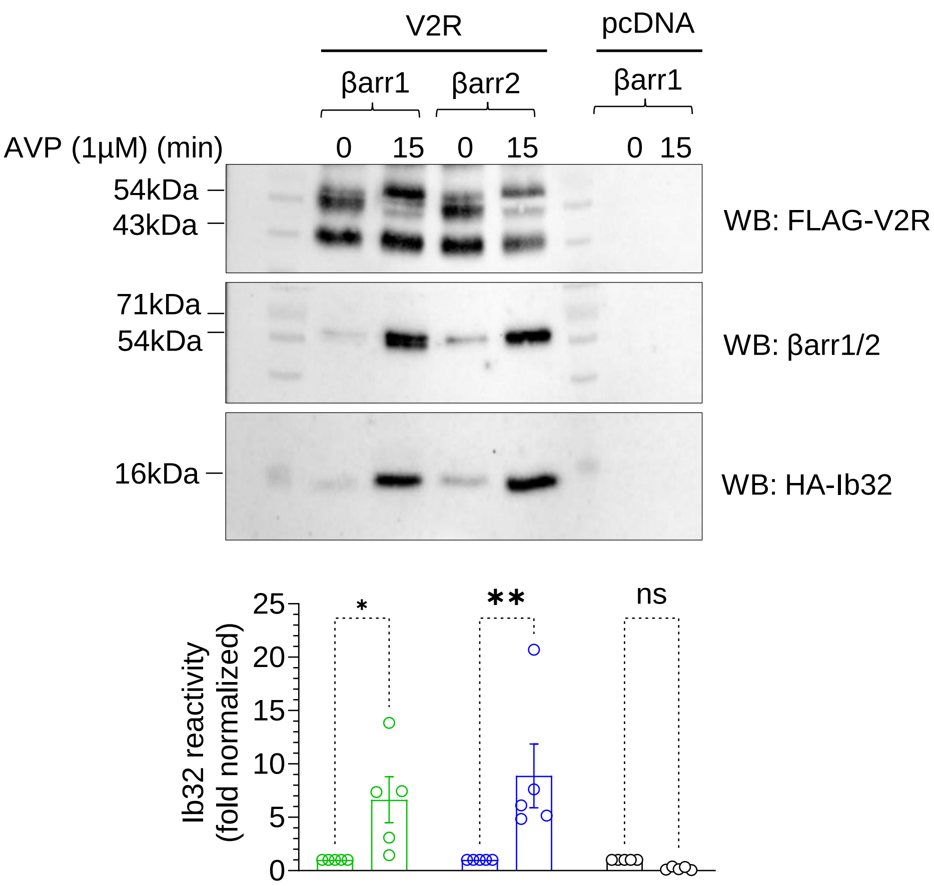


**Supplementary Figure 2. Ib32 reactivity measured using co-immunoprecipitation assay.** HEK-293 cells expressing the indicated constructs together with HA-tagged Ib32 were stimulated with AVP followed by pull-down using anti-Flag M1 antibody agarose and Western blotting. A rrepresentative blot (upper panel) and densitometry-based quantification (lower panel) are presented. The data represent mean±SEM fo five independent experiments, normalized with the unstimulated condition treated as 1, and analyzed using two-way ANOVA with Šídák's multiple comparisons test (**$p < 0.01$; *$p < 0.05$; ns, non-significant). The exact *p* values are as follows: pcDNA: 0 min vs. 15 min, 0.9758; V2R+βarr1: 0 min vs. 15 min, 0.0414; V2R+βarr2: 0 min vs. 15 min, 0.0033).

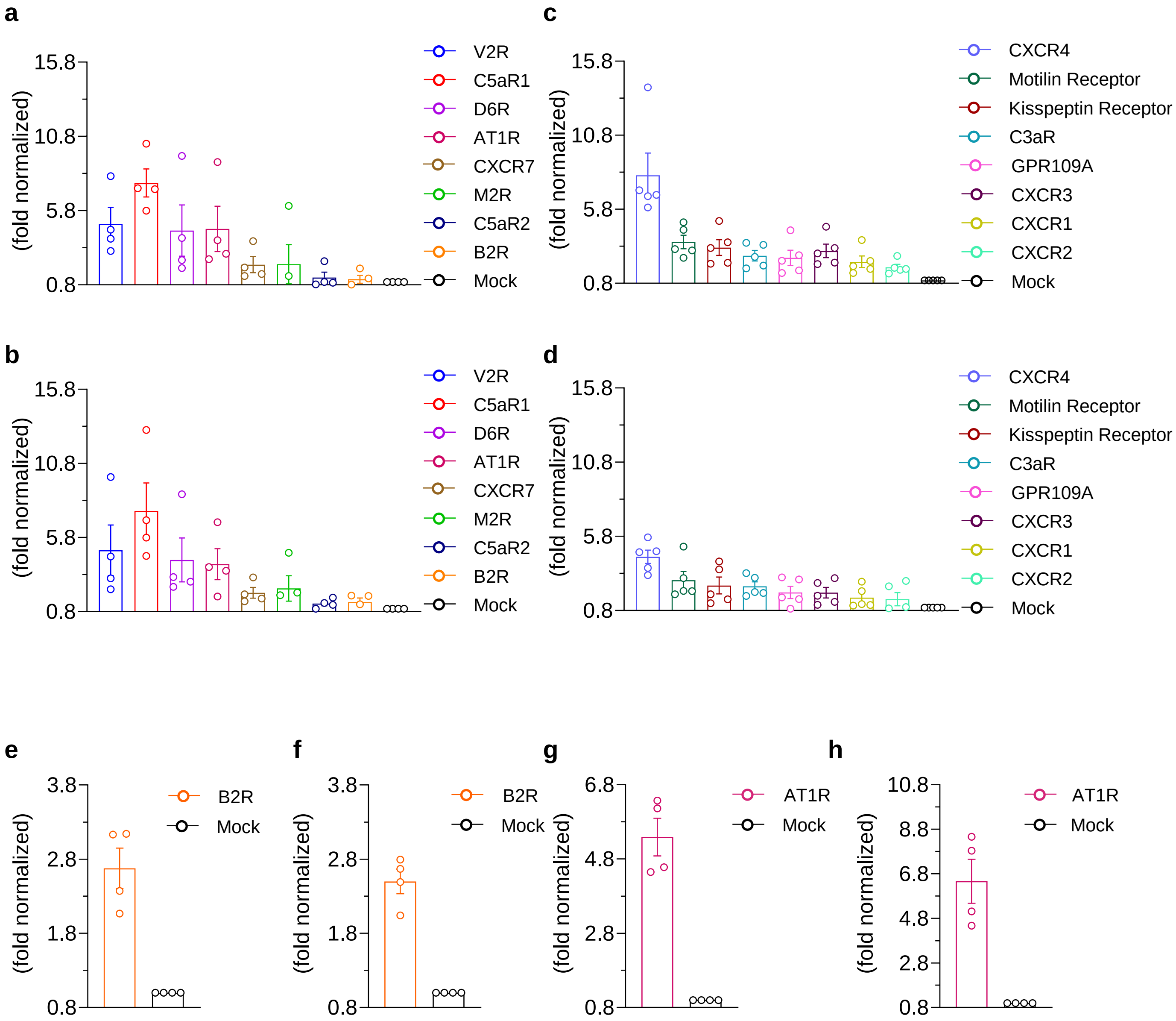


**Supplementary Figure 3. Surface expression of the indicated receptors. a-d,** Surface expression of the indicated receptors in the Ib32 reactivity assay data presented in Figure 2a, measured using whole cell ELISA. The data represent mean±SEM of 4-5 independent experiments, normalized as fold over mock-transfected condition. **e-h,** Surface expression of the B2R and AT1R in the Ib32 reactivity dose response data presented in Figure 2b-c, measured using whole cell ELISA. Data represent mean±SEM of four independent experiments, normalized as fold over mock-transfected condition.

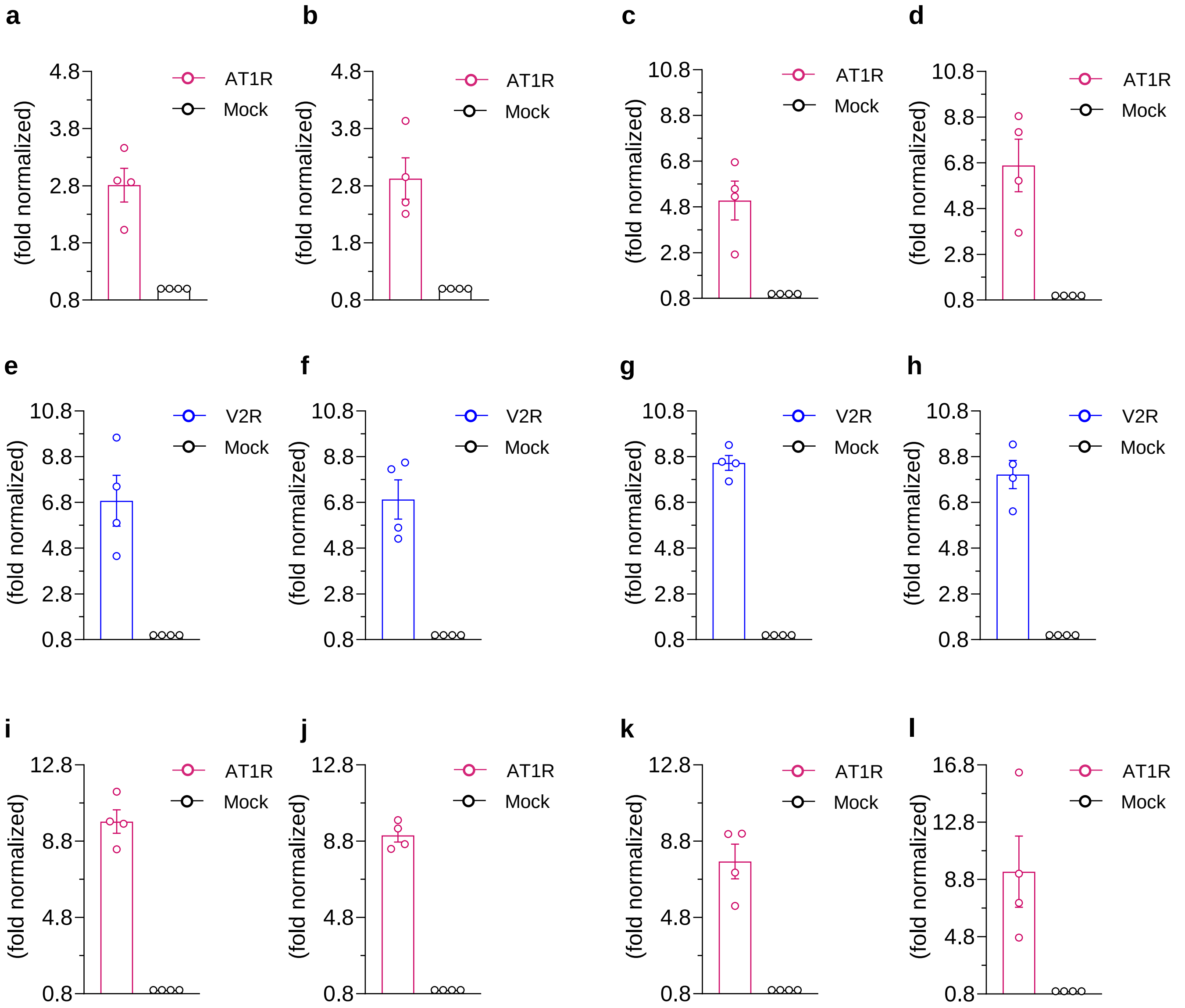


**Supplementary Figure 4. Surface expression of AT1R and V2R. a-d,** Surface expression of the AT1R in the Ib32 reactivity assay data presented in Figure 4a-d, measured using whole cell ELISA. Data represent mean±SEM of four independent experiments, normalized as fold over mock-transfected condition. **e-l,** Surface expression of the V2R and AT1R in the Ib32 reactivity assay data presented in Figure 5a-d, measured using whole cell ELISA. Data represent mean±SEM of four independent experiments, normalized as fold over mock-transfected condition.

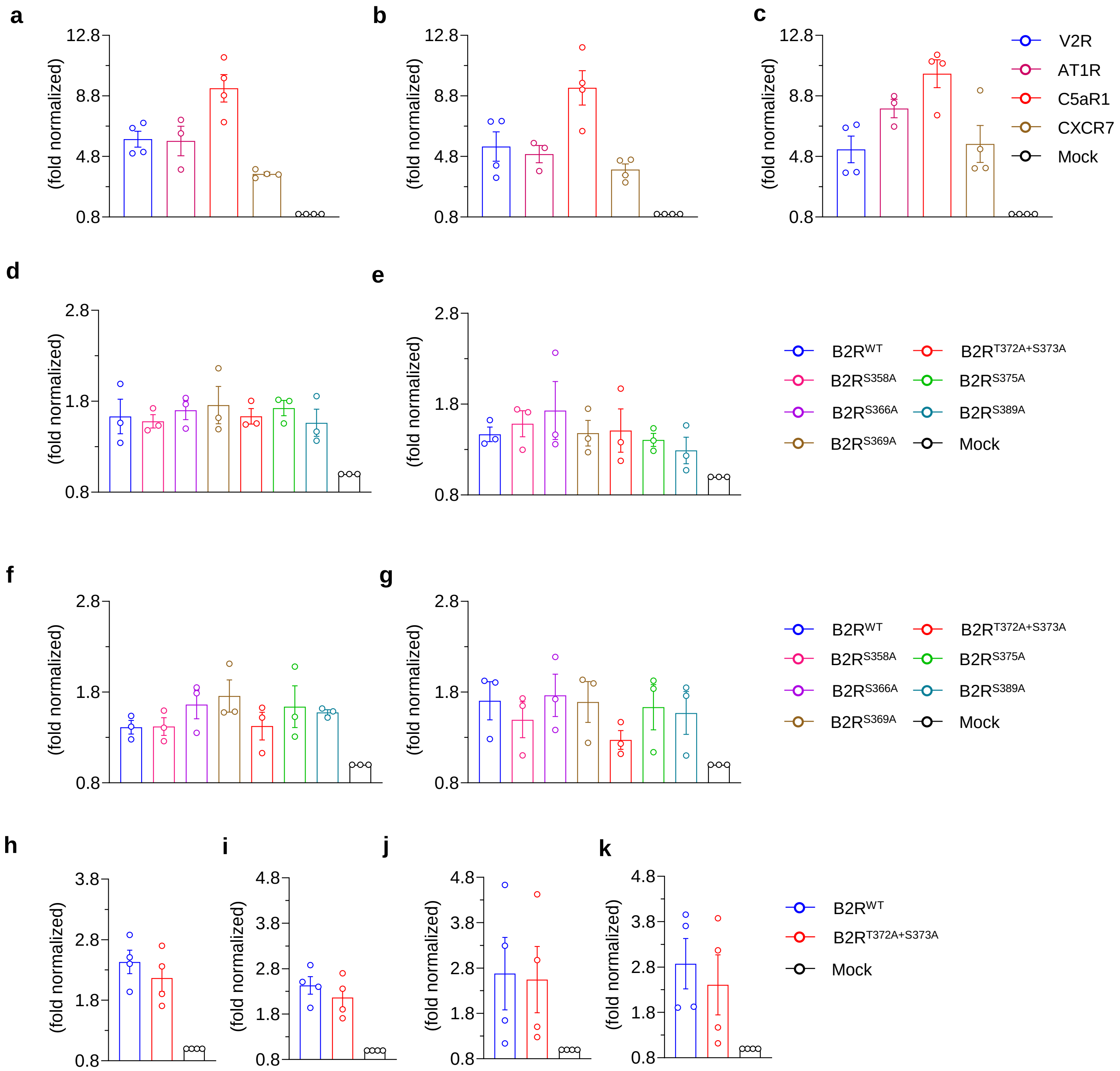


**Supplementary Figure 5. Surface expression of the indicated receptor constructs. a-c,** Surface expression of the indicated receptors in the Ib32 reactivity (a), Ib30 reactivity (b), and βarr recruitment (c) assay data presented in Figure 6a-c, measured using whole cell ELISA. Data represent mean±SEM of 3-4 independent experiments, normalized as fold over mock-transfected condition. **d-g,** Surface expression of the indicated B2R mutants in the βarr1/2 recruitment (d and f, respectively) and Ib32 reactivity assay for βarr1/2 (e and g, respectively) data presented in Figure 7b-c, measured using whole cell ELISA. Data represent mean±SEM of 3-4 independent experiments, normalized as fold over mock-transfected condition. **h-k,** Surface expression of the $B2R^{WT}$ and Thr372Ala+Ser373Ala mutant in the βarr1/2 recruitment (h and j, respectively) and Ib32 reactivity assay for βarr1/2 (i and k, respectively) data presented in Figure 7d-g, measured using whole cell ELISA. Data represent mean±SEM of 3-4 independent experiments, normalized as fold over mock-transfected condition.

| | |
|---|---|
| V2Rpp | : $^{343}$ARGRTPPSLGPQDESCTTASSSLAKDTSS$^{371}$ |
| C5aR1pp | : $^{331}$ESKSFTRSTVDTMAQKTQAV$^{350}$ |
| CXCR7pp | : $^{341}$ASRVSETEYSALEQSTK$^{362}$ |
| B2Rpp | : GGG-$^{363}$MENSMGTLRTSISVERQIHK$^{382}$ |
| AT1Rpp | : GGG-$^{321}$PPKAKSHSNLSTKMSTLSYRPSDNV$^{345}$ |

**Supplementary Figure 6. Sequence of the GPCR and ACR phosphopeptides used in the limited proteolysis assay.** The phosphorylated residues are highlighted in red color and the Gly-Gly-Gly sequence in B2Rpp and AT1Rpp indicates additional sequence designed for sortase ligation.
